## Supplemental data for "CNN explains tuning properties of anterior, but not middle, face-processing areas in macaque IT"

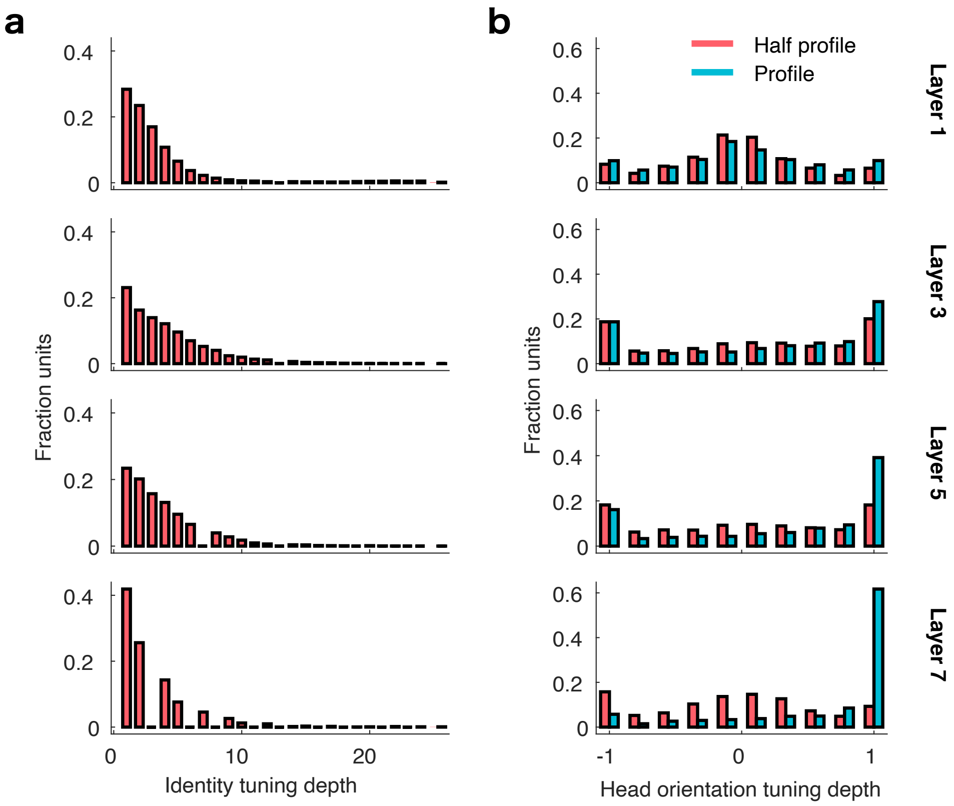


*Supplementary Figure 1: Additional results on identity selectivity and view invariance. (a) Identity selectivity. Identity tuning depth is defined as the index with half the maximum response when the responses to the 25 identities at the preferred head orientation were sorted. The distributions of identity tuning depths in most layers are more or less similar, where layer 7 gives a peakier distribution, somewhat compatible with AM, given in Figure 4G of the experimental study[3]. (b) View invariance. Head orientation tuning depth is defined as the ratio of the difference and sum of the average response to the frontal faces and that to the full (or half-profile) faces in the preferred direction. The distributions of head orientation tuning depths for full-profile faces are significantly different from experimental data in particular for the conspicuous peaks at -1 and 1, which is probably due to the weak view tolerance for full faces in AlexNet-Face (see Results). However, the distribution for half-profile faces in layer 7 has a slightly more pronounced bell-like shape as observed in AM, given in Figure 4H of the experimental study[3].*

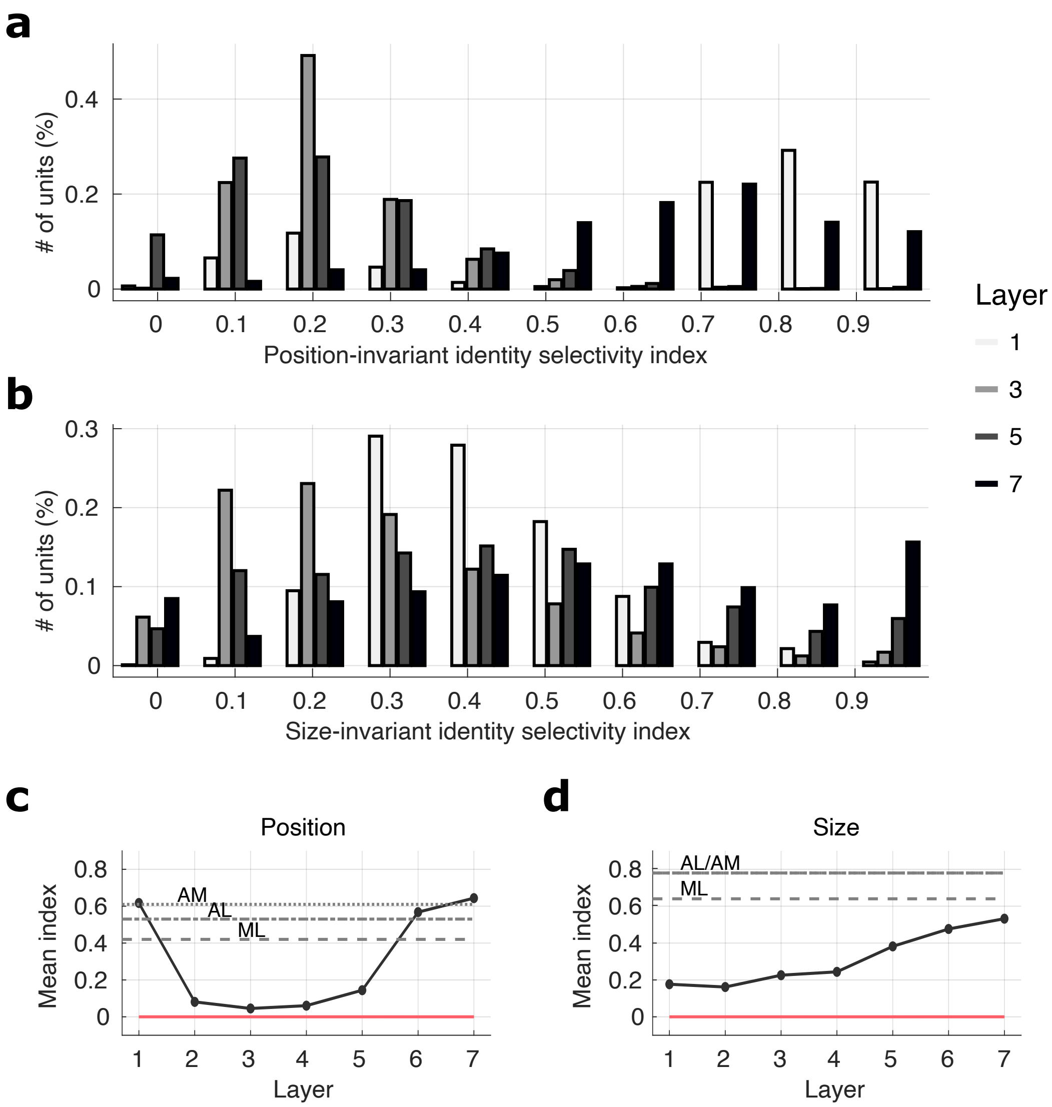


*Supplementary Figure 2: Additional results on position- and size-invariant identity selectivities. Position-invariant (or size-invariant) identity selectivity index is defined as the correlation coefficient between the population response vectors in the standard position (or size) and in the varied position (or size). For position variation, since the normal image size would exceed the boundary, we modified the protocol so that we used half-size images (112 pixels) and shifted the position upward, downward, leftward, or rightward by 28 or 56 pixels. (a) The distributions of position-invariant identity selectivity indices for different layers in AlexNet-Face model. (b) Analogous distributions of size-invariant identity selectivity indices. (c) The mean position-invariance indices across different layers in comparison with the corresponding experimental data on ML, AL, and AM[3]. Note that the comparison is only for reference due to the protocol modification. The spuriously high value in layer 1 is probably due to the spatial homogeneity of local textural features. (d) Analogous results on the mean size-invariance indices. In (c) and (d), the $\pm 2SD$ region of random cases (mean correlation coefficients between random response vectors drawn from Gaussian distribution) is highly concentrated to zero (red).*

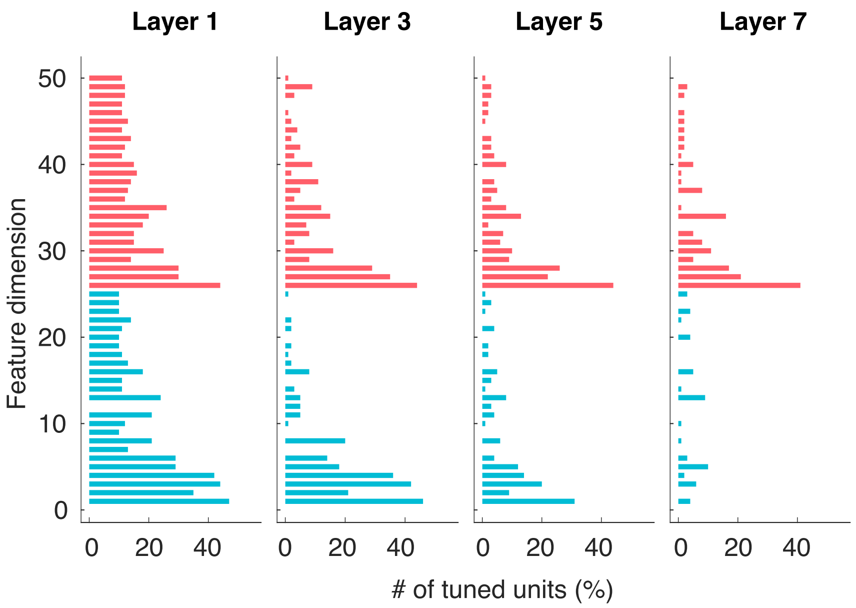


*Supplementary Figure 3: Additional results on shape-appearance preference from AlexNet-Face. The plots show the distribution of the number of significantly tuned units for each feature dimension (blue: shape, red appearance) in each layer. We used the same significance criterion as the experimental study[1]. Compare it with Figure 1H of the experimental study[1].*

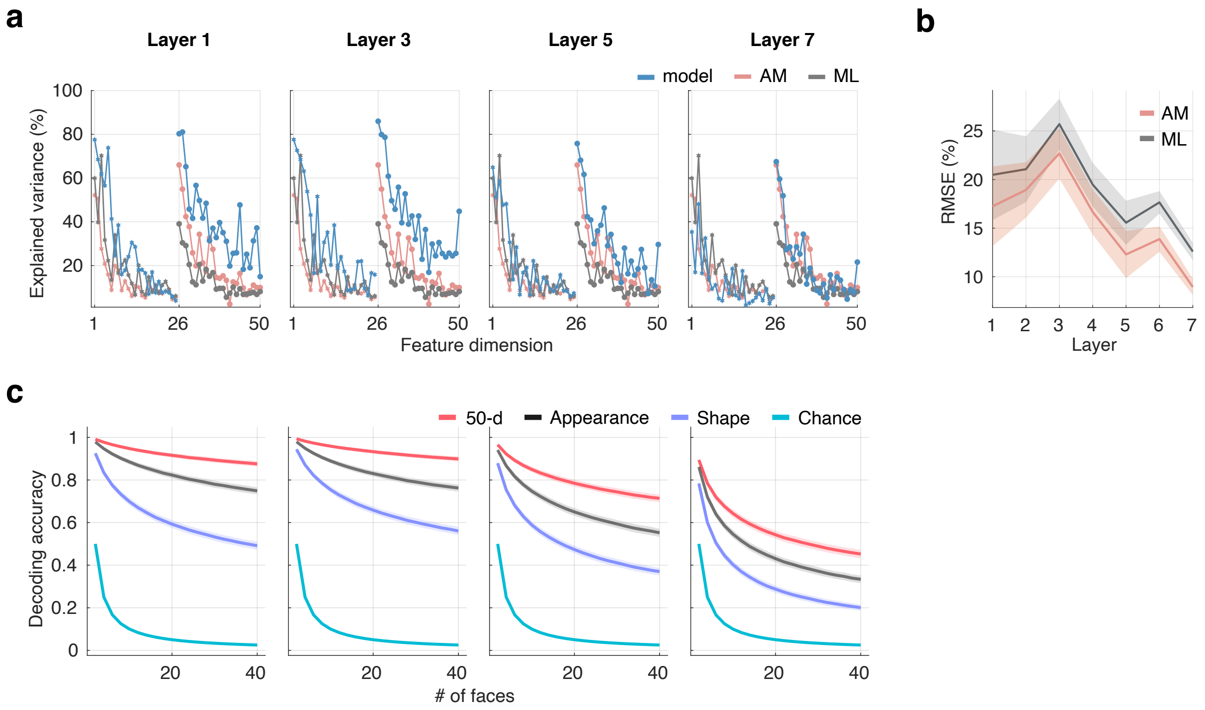


*Supplementary Figure 4: Additional results on decoding of facial features following the experimental study[1]. Since the size of unit population affects the performance of decoding, we randomly selected a subpopulation of 100 units from each layer, which is comparable to the size in the experiment. To decode a feature vector from the population responses for a chosen face image, we trained a linear regression model from the population responses to the remaining 1999 face images and predicted the feature vector for the chosen image. Using the leave-one-out approach, we continued this process for each of the 2000 images to decode the feature vectors for all images. (a) Decoding performance measured by explained variances. We calculated the ratio of explained variance in the actual feature values by the decoded feature values for each feature dimension (x-axis; 1–25: shape, 26–50: appearance). Each plot compares the results from a model layer (blue) with those from AM (red) and ML (gray), replotted from Figure 2C of the experimental study[1]. Note that all layers gave overall lower performance for shape dimensions than appearance dimensions, like AM and unlike ML. (b) Quantified comparison between the model layers and the experimental data. We calculated the root mean squared errors (RMSE) between the explained variances from each layer and from AM or ML. The shaded regions show the ±2SD range of the results from 100 cases with differently sampled 100-unit subpopulations. (c) Decoding performance measured by classification accuracy. We repeatedly sampled a subset of the face images and performed the nearest-neighbor algorithm on the decoded feature vectors against the actual feature vectors. Each plot shows the average decoding accuracy (red) as a function of the size of the subset of face images (2–40), along with the cases of decoding only the appearance dimensions (black) or the shape dimensions (purple), as well as the chance level (blue). The shaded regions indicate the standard deviation estimated using bootstrapping. Compare it with Figure 2D of the experimental study[1].*

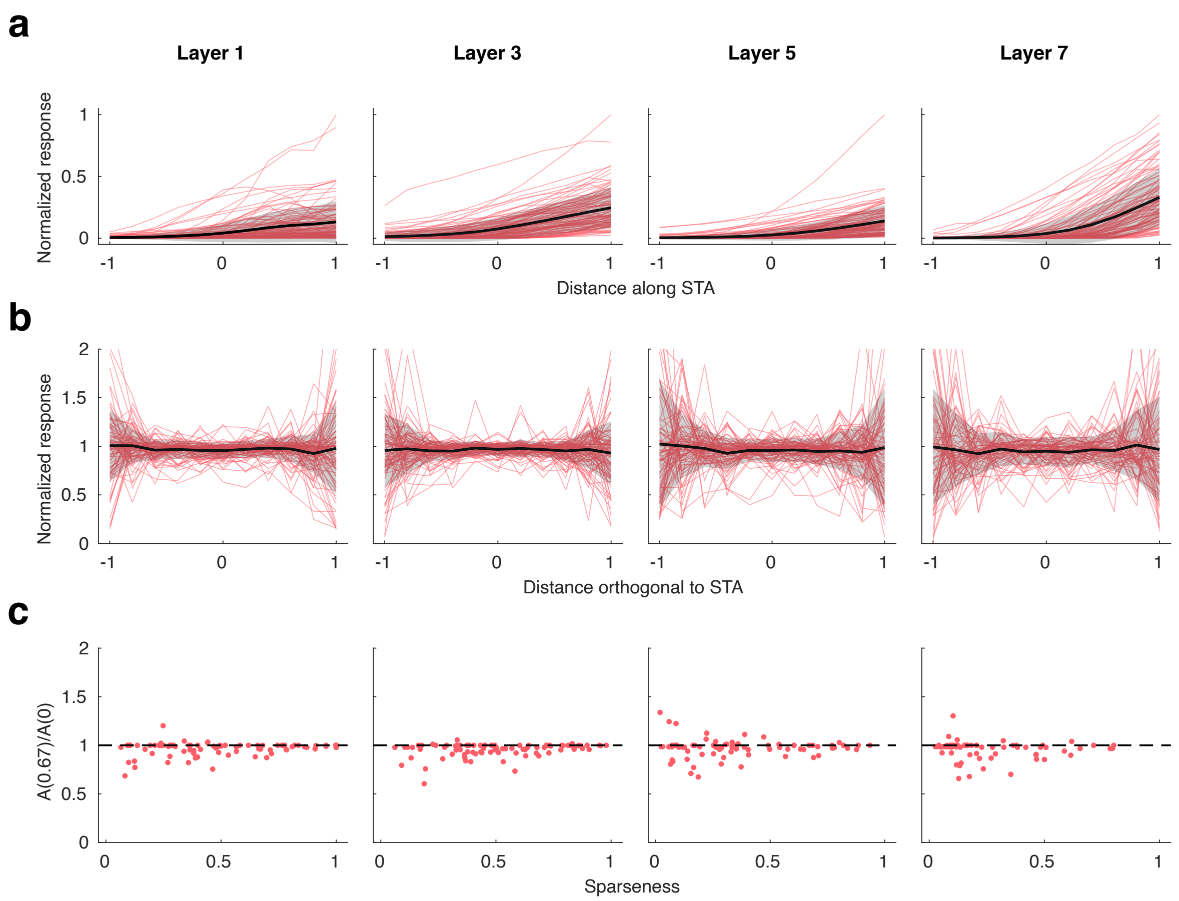


*Supplementary Figure 5: Additional results on ramp-flat tuning from AlexNet-Face, following the experimental method[1]. (a) Tuning curves along the STA axes for randomly selected 100 units (red), along with their mean (black) and s.d. (gray shade), in each layer. Each tuning curve was estimated by grouping the responses of the unit to the 2000 face images according to the distance between those faces and the mean face along the STA axis in the feature space. (b) Tuning curves along the principal orthogonal axes to STA (red), along with their mean (black) and s.d. (gray shade), in each layer. The principal orthogonal axis to a STA axis was obtained by first generating 2000 random feature vectors, next orthogonalizing these feature vectors with respect to the STA, and then taking the first principal component of these feature vectors. The tuning curve was estimated by similarly grouping the responses of the unit in the direction of the obtained orthogonal axis. After this, the tuning curve was fitted with a zero-centered Gaussian function ($k \cdot e^{-(x^2/\sigma^2)} + l$) and normalized by the value of the center of fit ($k + l$). We have not attempted the sparseness and noise matching unlike the experimental study[1] since the result (c) shows that the strength of nonlinearity seems mostly constant in our case. (c) The strength of nonlinearity of the tuning curve along the orthogonal axes to STA plotted against the sparseness: $(\sum_{i=1}^{N} R_i/N)^2/(\sum_{i=1}^{N} {R_i}^2/N)$, where $R_i$ are responses. The strength of nonlinearity was quantified by the ratio of the Gaussian fit at the surround ($x = 0.67$) and the center ($x = 0$).*

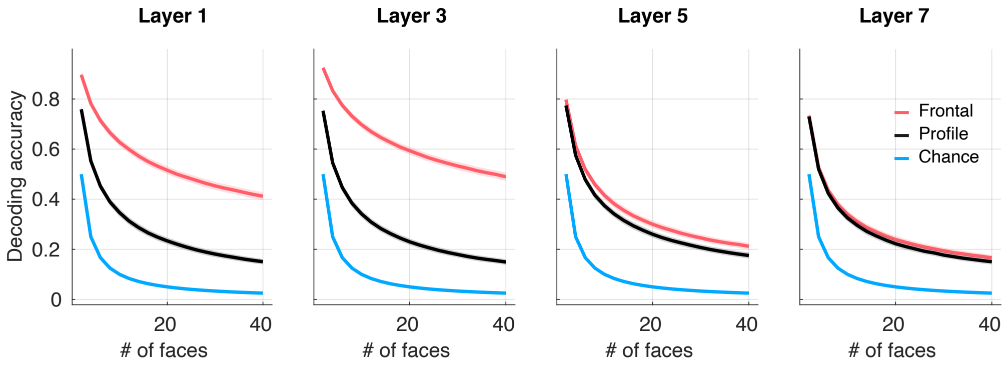


*Supplementary Figure 6: Additional results on view tolerance in the shape-appearance face space representation from AlexNet-Face. Plotted is the decoding accuracy for frontal faces (red) and profile faces (black), along with the chance level (blue), in each layer. Using the same method as in Supplementary Figure 4c, we trained each decoder for 46 units (similar to the experiment) using both frontal and profile faces[1].*

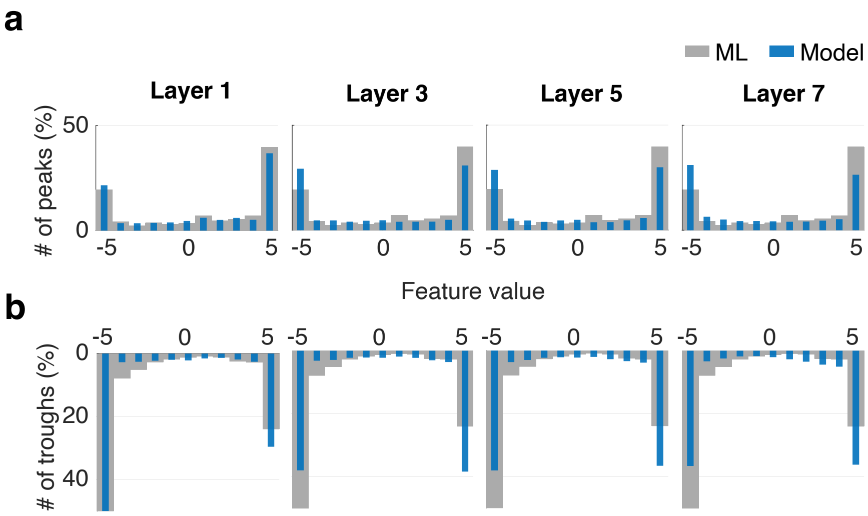


*Supplementary Figure 7: Additional results on facial geometry tuning in the cartoon face space from AlexNet-Face. Plotted is the distribution of the feature values that give the peak (a) or the trough (b) in each tuning curve from each model layer (blue), in comparison to the corresponding experimental data (gray), replotted from Figure 4 of the experimental study[2].*

| layer | AlexNet-Face | AF-5 | AF-6 | AF-8 | AF-9 | AF-h | AF-d |
|---|---|---|---|---|---|---|---|
| 1 | conv 11x11x3 | conv 11x11x3 | conv 11x11x3 | conv 11x11x3 | conv 11x11x3 | conv 11x11x3 | conv 11x11x3 |
| 2 | norm – pool – conv 5x5x96 | norm – pool – conv 5x5x96 | norm – pool – conv 5x5x96 | norm – pool – conv 5x5x96 | norm – pool – conv 5x5x96 | norm – pool – conv 5x5x48 | norm – pool – conv 5x5x192 |
| 3 | norm – pool – conv 3x3x256 | norm – pool – conv 3x3x256 | norm – pool – conv 3x3x256 | norm – pool – conv 3x3x256 | norm – pool – conv 3x3x256 | norm – pool – conv 3x3x128 | norm – pool – conv 3x3x512 |
| 4 | conv 3x3x384 | pool – fc 4096 | conv 3x3x384 | conv 3x3x384 | conv 3x3x384 | conv 3x3x192 | conv 3x3x768 |
| 5 | conv 3x3x384 | drop – fc 4096 | pool – fc 4096 | conv 3x3x384 | conv 3x3x384 | conv 3x3x192 | conv 3x3x768 |
| 6 | pool – fc 4096 | | drop – fc 4096 | conv 3x3x256 | conv 3x3x256 | pool – fc 2048 | pool – fc 8192 |
| 7 | drop – fc 4096 | | | pool – fc 4096 | conv 3x3x384 | drop – fc 2048 | drop – fc 8192 |
| 8 | | | | drop – fc 4096 | pool – fc 4096 | | |
| 9 | | | | | drop – fc 4096 | | |
| classif-cation accuracy | 72.78% | 65.96% | 68.07% | 69.04% | 68.04% | 64.15% | 70.08% |

*Supplementary Table 1: The architecture parameters of the trained CNN models used in this study. Each layer combines one or more processes of either convolutional filtering (conv), full connection (fc), channel-wise normalization (norm), max pooling (pool), or drop out (drop). The parameters for conv denote the width, height, and number of channels. The parameter for fc denote the number of units. The bottom row gives the classification accuracy of each model for held-out test data.*

| | VGG-Face | | AlexNet-Face | |
|---|---|---|---|---|
| No. | Layer | RF Size | RF Size | Layer |
| | relu1_1 | 3 | | |
| | relu1_2 | 5 | | |
| 1 | relu2_1 | 10 | 11 | relu1 |
| | relu2_2 | 14 | | |
| | relu3_1 | 24 | | |
| | relu3_2 | 32 | | |
| | relu3_3 | 40 | | |
| 2 | relu4_1 | 60 | 51 | relu2 |
| | relu4_2 | 76 | | |
| 3 | relu4_3 | 92 | 99 | relu3 |
| 4 | relu5_1 | 132 | 131 | relu4 |
| 5 | relu5_2 | 164 | 163 | relu5 |
| | relu5_3 | 196 | | |
| 6 | relu6 | 224 | 224 | relu6 |
| 7 | relu7 | 224 | 224 | relu7 |

*Supplementary Table 2: The layers of VGG-Face network that were selected for our analysis and the corresponding layers of AlexNet-Face network. Note the similarity between the indicated receptive field (RF) sizes of the corresponding layers.*
